# Fibroblasts drive differential response in immunotherapy via IL-6 in bioengineered 3D cancer constructs

**DOI:** 10.1101/2025.06.18.660473

**Authors:** Srija Chakraborty, Marco Rodriguez, Aleksander Skardal

**Affiliations:** Department of Biomedical Engineering, The Ohio State University, Columbus, Ohio, 43210; The Ohio State University Comprehensive Cancer Center, Columbus, Ohio, 43210; The Center for Cancer Engineering, The Ohio State University, Columbus, Ohio, 43210

**Keywords:** Cancer-associated fibroblasts, Natural killer cells, Chemokines, IL-6, Siltuximab

## Abstract

Despite various therapeutic advancements in cancer, heterogeneity in the tumor microenvironment is a major hindrance to the overall efficacy of treatment. In this study, we focus on the most abundantly found cellular stromal components – fibroblasts – which can be tumor-promoting or tumor-suppressing based on the extent of disease progression and numerous additional factors. In the studies described herein, we evaluated their role in influencing chemotherapy as well as in immunotherapy using colorectal cancer (CRC) cell line-derived 3D tumor constructs. Our initial efforts centered on common chemotherapeutic agents utilized in CRC. However, while the presence of fibroblasts often did influence chemotherapy efficacy, a clear and consistent increase or decrease in efficacy was not obvious. We then focused subsequent studies on natural killer (NK) cell-based immunotherapy efficacy. There are numerous components and pathways involved in tumorigenesis and tumor-immune cell interactions, and the crosstalk between fibroblasts and immune cells like NK cells is still not well-defined. Fibroblasts have been documented to be a major player with respect to immune modulation and understanding this in greater detail can elucidate more options for anti-tumor immunity. Specifically, we used an extracellular matrix-based hydrogel platform to generate 3D CRC constructs to evaluate how fibroblasts affect tumor cell viability in presence and absence of NK cells. In general, fibroblasts aid the tumor cells in our models. We then identify a cytokine-based pathway (interleukin 6 [IL-6]) via which ‘activated’ fibroblasts engage with the tumor cells and show how NK cell effector function can be restored when the IL-6 pathway is blocked in a melanoma-based tumor-on-a-chip platform.

**Graphical abstract:** 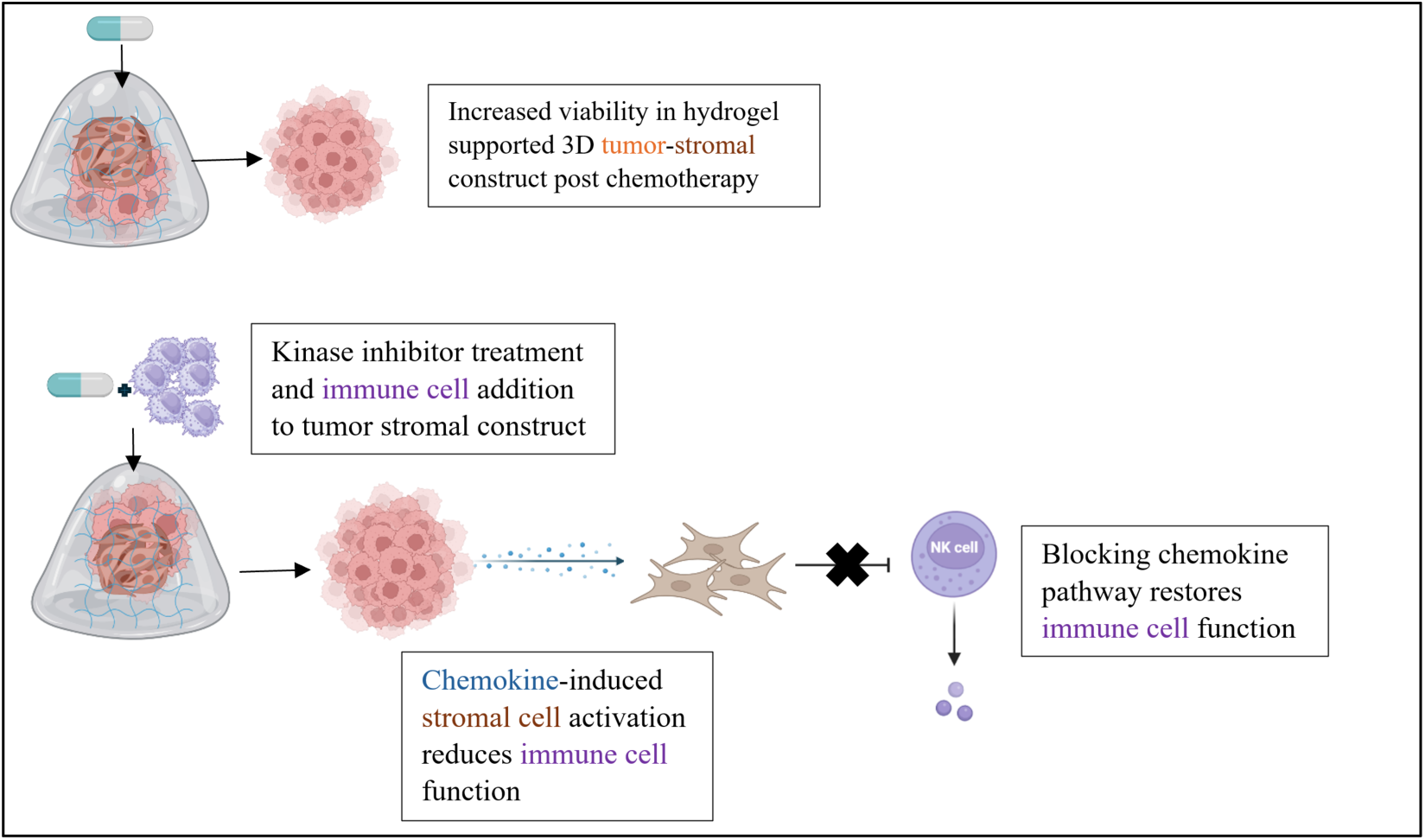

## 1. Introduction

Colorectal cancer (CRC) or colon/bowel/rectal cancer-affects the colon and rectum. It can develop from polyps/outgrowths lining the colon and gradually turn cancerous without treatment. According to the World Health Organization, colon cancer is the second leading cause of cancer-related deaths worldwide. There were an estimated 106,970 new cases of colon cancer in 2023 with 52,550 projected deaths from colorectal cancer (including both colon and rectal cancers).^1^ Colon cancer has a higher chance to metastasize to distant organs once it has reached the lymph system. The American Joint Committee on Cancer defines ‘surgical staging’ for CRC based on the ‘TNM’ criteria which denotes extent or size of tumor (T), the spread to nearby lymph nodes (N) and the spread or metastasis to distant sites (M). Thus, the 5-year survival rate for colorectal cancer varies depending on the stage at which it is diagnosed with ‘stage 0’ being where cancer cells are limited to the innermost lining of the colon and ‘distant or stage IV’ being where the cancer has metastasized to distant organs, such as the liver or lungs, the five-year survival rate significantly decreases to about 14%.^2^ Treatment can be combinational with surgery, chemotherapy, and even radiation therapy. Chemotherapy can be neoadjuvant or adjuvant and can be administered to the patient systemically or locally. Some commonly used drugs (as monotherapy or in combination) for CRC include 5-Fluorouracil, Oxiplatin, and Irinotecan.^3,4^ More recently, targeted therapies and immunotherapies have also been proven to be effective. However, as is with other types of cancer, patient heterogeneity is a major factor in deciding the extent of therapeutic success.

There are several methods and models which have been used to study colorectal cancer in the laboratory. Two-dimensional (2D) culture methods, despite their low cost and maintenance, easy reproducibility of results and standardized techniques, can neither faithfully recapitulate complexity of the CRC tumor microenvironment (TME), nor can effectively evaluate involvement of non-tumor cell types. Hence, 3D *in vitro* tumor models are being increasingly used to model and study the different aspects of CRC and other cancers. These models have also become more complex with time, starting from the hanging drop method to generate simple spheroids to the more sophisticated organoid self-assembly model.^5^ CRC cell aggregates or spheroids represent a 3D avascular model that can mimic cell-cell interactions, but often ignore cell-matrix interactions. Extracellular matrix (ECM)-derived hydrogels offer alternative methods to model the 3D tumor biology, both in terms of cell-cell and cell-matrix interactions. Collagen is the most abundant protein present in our extracellular matrix (ECM), and it is also present in the TME. Collagen type I hydrogels are often used in the development of 3D models of CRC due to their ability to provide an ECM-like structure. However, collagen is not the only ECM component in most tissues and tumors. Hyaluronic acid (HA) is one of the next most prevalent ECM components in most tumors, and part of a collagen-HA hydrogel platform developed and deployed in previous studies by our team. These hydrogels-made of natural components that are synthetically modified to support covalent crosslinking on demand. The subsequent multi-component hydrogels enable the study of cell–cell and cell–matrix interactions as well as tumor cell invasion.^5^

Bioengineered tissue and tumor models can be further advanced using microfluidic device technologies, resulting in organ- or tumor-on-a-chip systems. Our lab and others have deployed such platforms to assess interactions between multiple tissue types following drug insults, and to study complex disease states, such as metastasis from one primary site to one or more downstream target sites.^6–10^ These microfluidic devices or tumor-on-a-chip platforms perfuse cell culture medium through the cellular structures enabling biochemical stimuli like oxygen and nutrient diffusion. Thus, these systems can be used to study pharmacokinetics and drug delivery in great details.^11^

As with other malignancies, the TME in CRC is also heavily heterogeneous and different components, cellular – including stromal and immune cells – or acellular – including ECM and cytokines – have different contributions towards aiding and/or suppressing tumor formation and progression. Stromal residents like fibroblasts and more specifically cancer-associated fibroblasts, or CAFs, have various roles in aiding CRC cells through direct secretion of soluble factors that upregulate or downregulate other pathways.^12,13^ CAFs have been shown to secrete IL-6 which can upregulate Leucine Rich Alpha-2-Glycoprotein 1 (LRG1) expression in colorectal cancer and is also correlated with prominent CAF marker alpha smooth muscle actin (α-SMA) and eventually with metastasis and thus poor prognosis.^14,15^

Natural killer (NK) cells, among other innate immune cells, are often participants of post-inflammatory cancer mechanisms. Studies in CRC show that during tumor progression, inflammation is commonly observed, resulting in infiltration of NK cells and an increased presence at tumor sites as part of the immune system’s efforts to curtail tumor progression. Through a cascade of events including cytokine secretion, cytokines and hormones such as leptin can trigger M1 macrophages to secrete IL-1β which in turn can result in lymphocyte cell-mediated production of pro-inflammatory IL-6, which depresses NK cell effector function.^16^ While NK cells can be part of a natural response to limit tumor progression, they can also be harnessed as a form of cellular immunotherapy.^17^ Cytokine-centric therapeutic approaches (IL-2 and IL-15) can promote NK cell activation, and expansion, have been tested in several preclinical and clinical settings. However, the use of both cytokines has shown adverse effects on patients as well.^18^ Alternatively, NK cells can be isolated from a patient, expanded, and returned to the patient as a therapy. Additionally, the NK-92 NK cell line has also been explored in the clinic.^19,20^ In the TME fibroblasts or more specifically cancer-associated fibroblasts, can suppress the function of NK cells in colorectal cancer in various ways. CAFs can remodel the ECM and secrete ECM proteins in the tumor stroma, which can block and thus reduce efficacy of NK cells. Some immune cells secrete cytokines or chemokines to activate normal/resting fibroblasts to CAFs promoting the formation of physical barriers in the TME, reducing the physical contact between immune cells and cancer cells, and therefore the immune surveillance of tumor cells. CAFs can also inhibit the expression of perforin and granzyme B but also reduce the expression of tumor necrosis factor-α (TNF-α) and interferon gamma (IFN-γ).^21–23^

In the studies proposed herein, we used bioengineered CRC tumor constructs supported by our team’s collagen-HA ECM hydrogel system to assess the influence of fibroblasts on chemotherapies and NK cell-based immunotherapy. Notably, based on the results described below, our attention shifted to focus on the NK cell immunotherapy. Our hypothesis was that the addition of fibroblasts affects chemotherapy and immunotherapy response in hydrogel-supported tumor constructs (TCs) constructed with CRC cell lines. This response is directed by extent of aggression of the tumor cells. Moreover, presence of fibroblasts dictates effector function of immune cells via specific pathways. Hence, this study has two major sections-chemotherapy-based and immunotherapy-based (**as schematically represented in the graphical abstract**).

## 2. Results

### 2.1 Fibroblasts show differential outcomes in colorectal TCs post chemotherapy

We initially wanted to query the influence of fibroblasts on chemotherapeutic efficacy using our collagen-HA hydrogel-supported tumor constructs created with a panel of CRC cell lines that we have deployed in previous studies.^7,24–26^ In these studies, we evaluated recovery response of tumor cells post treatment with these chemotherapeutic drugs. 5-FU resembles the pyrimidine molecule in DNA and RNA and in the patient’s body, incorporates itself into the DNA or RNA of cancer cells to inhibit thymidylate synthase, needed for DNA replication and repair, thus preventing tumor cell proliferation. Intravenous administration of 5-fluorouracil (5-FU) was the treatment choice for advanced CRC for a number of years, but current options include combinations of 5-FU with cytotoxic agents such as oxaliplatin and irinotecan, resulting in the chemotherapeutic regimens FOLFOX (5-FU + leucovorin + oxaliplatin), FOLFIRI (5-FU + leucovorin + irinotecan), and FOLFOXIRI (5-FU + leucovorin + oxaliplatin + irinotecan).^27^ In our studies, CRC tumor constructs (TCs) were formed by encapsulating CRC cells (Caco-2, SW480, or HCT-116) with or without fibroblasts (human normal lung fibroblasts [HNLF]). The most discernable results were seen with 5-FU treatment. Post 5-FU treatment, Caco-2-HNLF samples show higher viability than only Caco-2 TCs, at higher drug concentrations and across later time points in both 2D culture as well as 3D TCs (**Fig 1i**). For the moderately aggressive or epithelial SW480 TCs, the viability is somewhat similar in the 2D samples made with and without fibroblasts (**Fig 1ii**). However, for the 3D TCs, cell density is increased, and viability is higher in the TCs containing fibroblasts (**Fig 1iii**). For the control groups, Caco-2 TCs (**Fig 1 iv**) show cells dying off in the center, which could be due to the growth media being more accessible to peripheral regions. SW480 TCs show robust cell growth, and for the HCT-116 TCs, there are cell clusters forming within the constructs as well, indicating high cellular viability.

**Figure 1:**
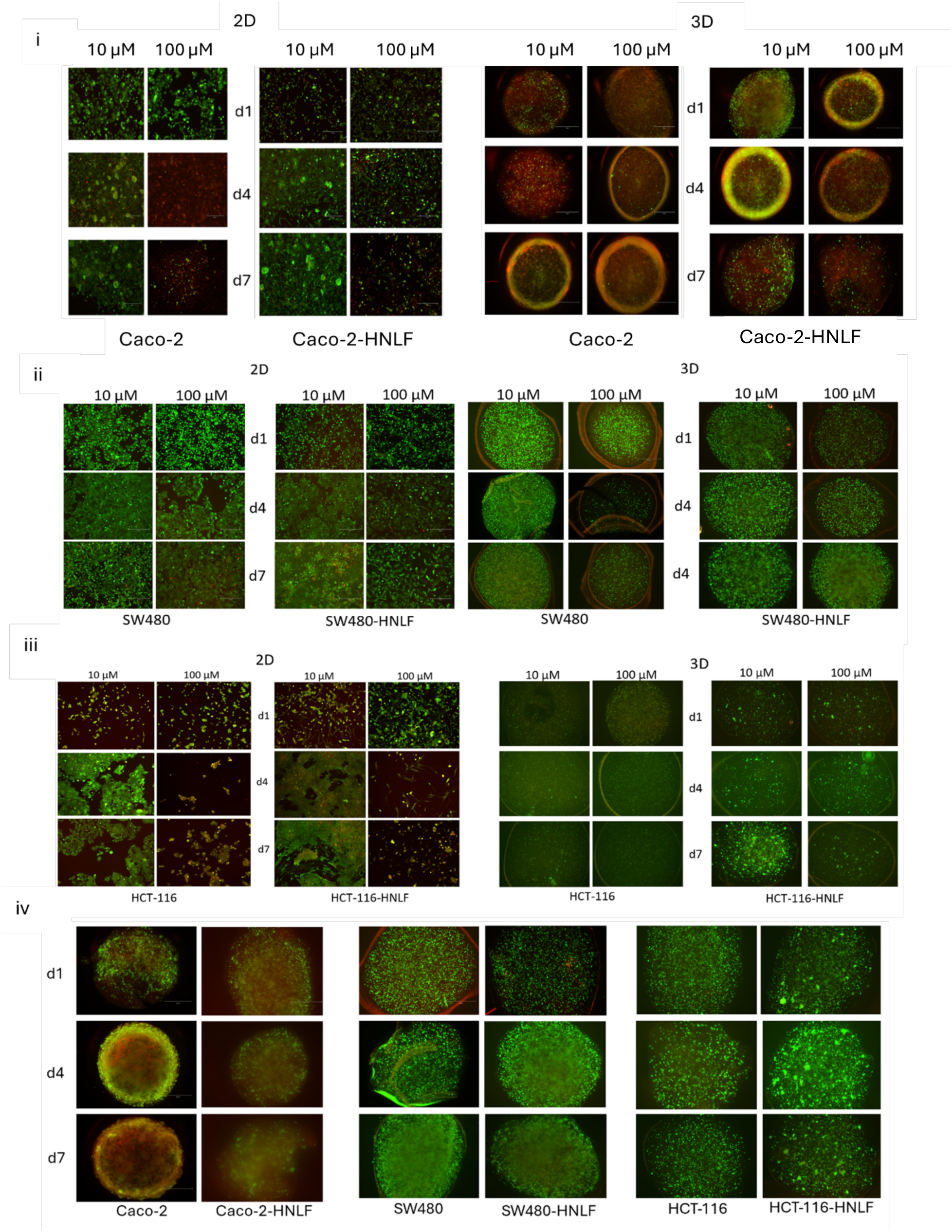
Post 5-FU treatment-viability in 2D tissue culture and 3D hydrogel-supported colorectal cancer tumor constructs made with and without fibroblasts via LIVE/DEAD assay on days 1,4, and 7. (i) Representative imaging of LIVE/DEAD viability assay in Caco-2 cells with and without fibroblasts in 2D culture and 3D tumor constructs showed higher viability in 100 μM treated samples in both 2D and 3D types. (ii) There was not as much difference in viability between tumor cell only and tumor cell-fibroblast samples for SW480 cells, in both 2D and 3D conditions, although the cellular growth is visibly denser than Caco-2 samples. (iii) Viability is higher HCT-116-HNLF tumor constructs compared to HCT-116 only samples, although this difference in viability is not as clear for the 2D samples. (iv) No drug-treated control samples only for TCs for all three cell lines, with and without fibroblasts showed increasing viability with increasing degree of aggressiveness from Caco-2 to HCT-116 samples. Scale bar - 300 μm. Green—calcein AM-stained viable cells; Red—ethidium homodimer-1-stained dead cell nuclei.

Other studies were done with Cisplatin and Regorafenib where viability was measured via the same assays after treatment with these drugs (mentioned in the supplementary section, **Fig S1 and S2**). However, the response was not uniform for all the conditions tested and how fibroblasts influenced treatment was not nearly as clear nor consistent as we saw for the TCs that received 5-FU-based therapies. Likewise, our team often pairs LIVE/DEAD staining with ATP quantification for stronger corroboration of our data sets. As shown in **Fig S3, S4, and S5**, there is little correlation between LIVE/DEAD staining and ATP quantification. This could very likely be because the mechanisms of actions between these chemotherapies are all quite different. As a result of these data, because our team also has an interest in evaluating how components of the TME influence immunotherapies, we added a focus on the influence of fibroblasts on cellular immunotherapy. Specifically, we performed a next set of studies in which we evaluated NK cell-based tumor cell killing with and without the presence of fibroblasts, which became our primary focus.

### 2.2 Kinase inhibitor-guided NK cell effector function is most prominent in least aggressive colorectal cancer TCs, and reduced NK cell effector function is seen in TCs in presence of fibroblasts

Kinase inhibitor treatment has been used in the treatment of solid tumors. These are increasingly being used due to their ability to target specific enzymes involved in the regulation of cell signaling pathways. These inhibitors work by blocking the activity of kinases, which are often upregulated in cancer cells and contribute to tumor growth, survival, as well as metastasis. Thus, small molecule inhibitors can interfere with key processes like angiogenesis, cell cycle progression, and apoptosis evasion. One such drug is Alisertib (ALS), currently being investigated as a potential treatment for melanoma (although in combination with other therapies like trametinib), as it targets Aurora kinase A, a protein that plays a role in cancer cell division. ALS is also documented to render tumor cells senescent and help in reducing their proliferation.^28^ Abemaciclib is another kinase inhibitor which targets the CDK4/6 pathway and has been used in breast cancer studies as well as melanoma cases.^29,30^

Also, upon treatment with these drugs, many tumor cells can be driven into senescence. Once rendered senescent, the tumor cells often secrete a shifted cytokine and chemokine profile, including increases in chemokines such as CCL5 and CXCL10 which we hypothesized can attract the NK cells towards the TCs and hence augment their cytotoxicity and tumor elimination. Therefore, to evaluate how tumor cells behave under different treatment conditions, we used our CRC cells and treated them with ALS and ABE followed by NK cells, to boost immunotherapy efficacy. After that, we added in fibroblasts to our CRC TCs and evaluated viability as well as NK cell effector function. These studies (unless otherwise mentioned) were done using well-plates where the TCs were made with CRC cells, with and without the addition of fibroblasts. The NK cells were introduced to these constructs via transwells (**Fig 2i**) through which they would have to infiltrate down to the TCs. NK-92 addition to small molecule kinase inhibitor-treated CRC only TCs show greater cytotoxic effect on least metastatic samples i.e. Caco-2 (**Fig 2ii**). Viability is lowered in ABE-treated samples as shown via the ATP assay (**Fig 2iii**). NK-92 effector function is confirmed via the presence of Granzyme B as well as TNF-α and IFN-γ (**Fig 2iv-vi**). Here, NK cells carry out their effector functions more prominently in drug-treated samples, specifically Granzyme B and TNF-α for ALS-treatment. Overall, ALS treatment shows better outcomes than ABE. With respect to TCs made with both Caco-2 and fibroblasts, significant difference in viability is not visible between samples not treated with NK cells and those where NK cells were added, as represented by the LIVE/DEAD images (**Fig 3i-ii**). However, the ATP assay showed highest viability in the control groups compared to the drug-treated groups. We attributed this discrepancy to the fibroblasts lowering NK cell cytotoxicity. (**Fig 3iii**). Additionally, here, NK cell effector function was confirmed via Granzyme B, TNF-α and IFN-γ-based ELISAs (**Fig 3iv-vi**). Granzyme B and TNF-α are upregulated more in ALS-treated samples compared to the control groups while for IFN-γ this upregulation is not as significant.

**Figure 2:**
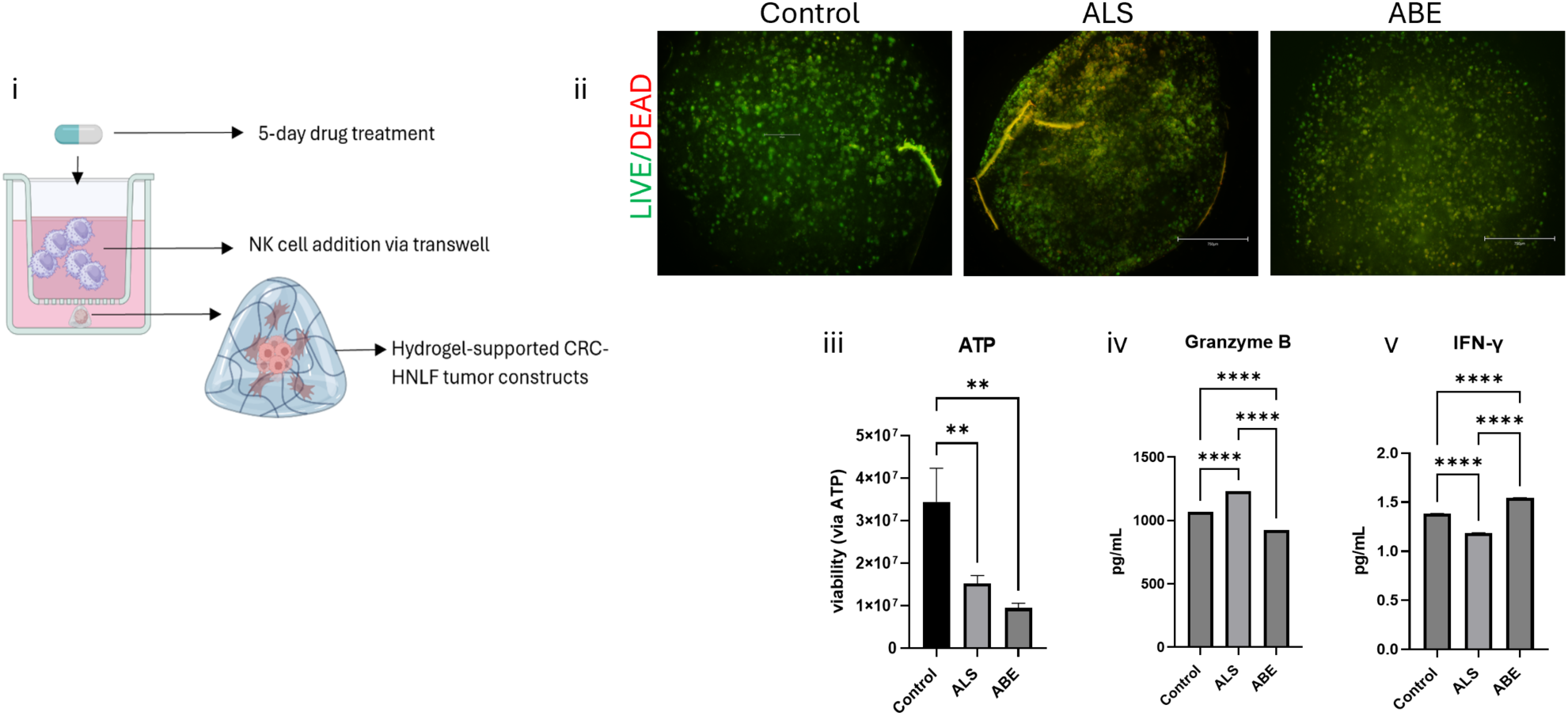
Schematic of tumor cell-fibroblast construction, drug treatment and NK cell addition as well as viability and NK cell function in tumor-only constructs. (i) Caco-2 colorectal cancer (CRC) cells are used with human normal lung fibroblasts (HNLF) to make tumor constructs supported with extracellular matrix-based collagen-hyaluronic acid hydrogels. NK-92 cells are added via transwells after 5 days of kinase inhibitor drug treatment. (ii) Representative imaging of LIVE/DEAD viability assay in Caco-2 only TCs with NK cell treatment showed greater viability in control image (iii) ATP assay confirmed lower viability in drug-treated specially ABE-treated samples (iv-vi) ELISA results validated NK cell effector function and increased function in ALS-treated samples compared to controls mainly for Granzyme B and TNF-α. Scale bar - 750 μm. Statistical significance: * p < 0.05, ** p < 0.01, *** p < 0.001, **** p < 0.0001. Data are represented as mean + SD. Green - calcein AM-stained viable cells; Red - ethidium homodimer-1-stained dead cell nuclei.

**Figure 3:**
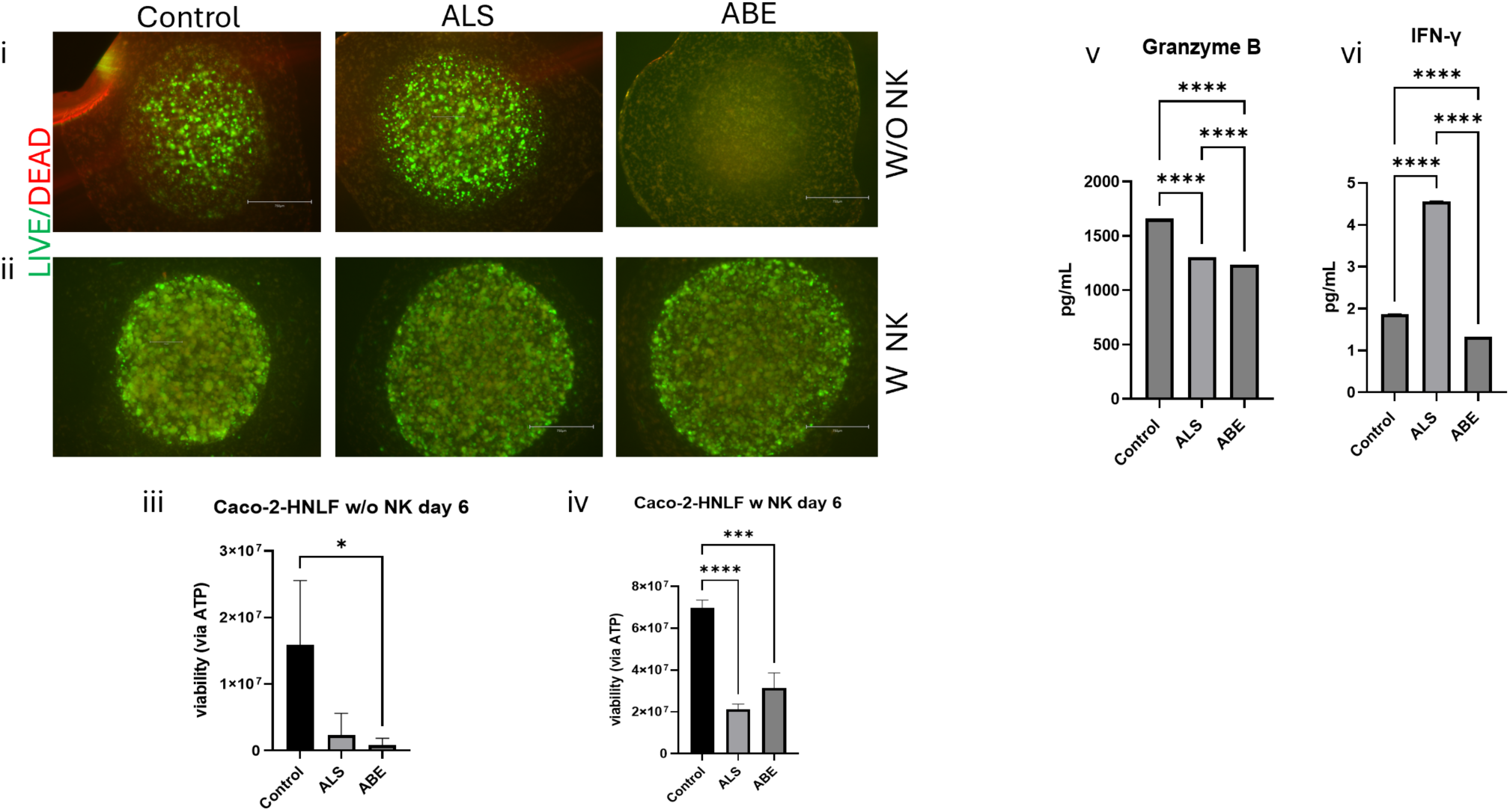
Viability assay and NK cell function in tumor cell-fibroblast constructs, with and without NK cell treatment. (i) Representative images of LIVE/DEAD viability assay showed higher viability in samples without NK cell addition compared to (ii) those with NK cell treatment (iii-iv) ATP assay showed higher viability in samples without NK cell addition (v-vii) ELISA results validated NK cell effector function and increased function in ALS-treated samples compared to controls for Granzyme B, TNF-α and IFN-γ. Scale bar - 750 μm. Statistical significance: * p < 0.05, ** p < 0.01, *** p < 0.001, **** p < 0.0001. Data are represented as mean + SD. Green - calcein AM-stained viable cells; Red - ethidium homodimer-1-stained dead cell nuclei.

### 2.3 Expression of E-cadherin is lowered in tumor-stromal TCs while expression of IL-6 is increased

The loss of E-cadherin expression is usually a marker for tumor progression. This has also been associated with loss of differentiation in CRC, although expression levels of β-catenin have also been used in conjunction.^31^ Moreover, reduced E-cadherin expression can lead to activation of fibroblasts resulting in the formation of CAFs.^32^ In our Caco-2 samples, there is reduced E-cadherin expression which points towards tumor aggression and resistance to chemotherapy (**Fig 4i**). Some non-specific fluorescent signal is observed, but E-cadherin most robustly present in Caco-2-only samples. The chemokine profiles were similar for the samples pertaining to the SW480 and HCT-116 based constructs (**Fig 4iii and v**), but E-cadherin staining not being conclusive we proceeded with Caco-2 samples only. Additionally, ELISA panels for markers of ‘activated’ fibroblasts and/or fibroblast involvement showed significantly increased presence of IL-6, IL-8, CCL5, and CXCL10 in the tumor cell-stromal cell co-cultures compared to both tumor cell only and stromal cell only constructs for Caco-2, Sw480, and HCT-116 samples (**Fig 4ii, iv, and vi respectively**). CCL5 has been shown to aid tumor growth via fibroblast recruitment. CCL5 derived from CAFs can also facilitate metastasis in hepatocellular carcinoma. CXCL10 on the other hand can have dual roles in cancer, it can enable tumorigenesis, but also inhibit tumor growth.^33–35^ Although this upregulation in the different chemokines was uniformly observed for all three colorectal cancer cell lines used, it was most prominent in Caco-2 TCs.

**Figure 4:**
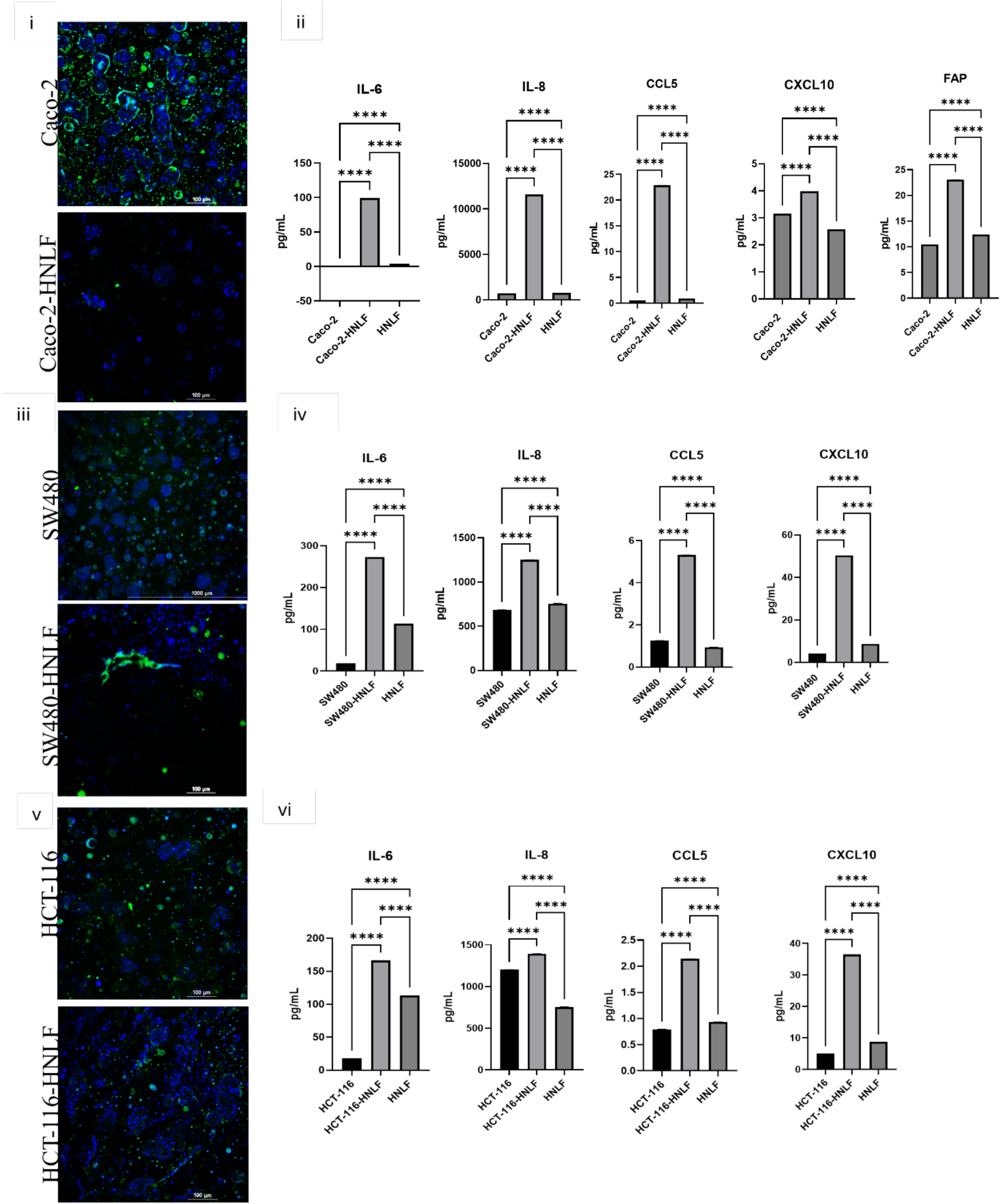
E-cadherin expression and chemokine profile assessment in tumor cell, tumor cell-fibroblast and fibroblast only constructs. (i-top) Caco-2 construct showed higher expression of E-cadherin compared to (i-bottom) Caco-2-HNLF construct. (ii) ELISAs done on Caco-2HNLF constructs also showed increased expression of several chemokines like IL-6, IL-8, CCL5 and CXCL10. FAP-ELISA done to validate activated state of fibroblasts showed increased expression in Caco-2-HNLF samples compared to Caco-2 only and HNLF-only samples. (iii-top and bottom) E-cadherin expression was not clear in either sample although (iv) chemokine profiles were like those seen for Caco-2, with and without HNLF groups. (v-top and bottom) E-cadherin expression was not clear in either sample although (vi) chemokine profiles were like those seen for Caco-2, with and without HNLF groups. Hence FAP ELISA was not done for either SW480 or HCT-116 containing samples. Scale bars - 100 μm. Statistical significance: * p < 0.05, ** p < 0.01, *** p < 0.001, **** p < 0.0001. Data are represented as mean +SD. Green - E-cadherin; Blue - DAPI-nuclear stain.

### 2.4 Increasing dosage of IL-6 blocker in TCs improves NK cell function on tumor cell viability

CAFs are known to interfere with E-cadherin expression via TGF-β (and later EMT), IL-6, and other cytokines. IL-6 and IL-8 cytokines are known to be indicators of fibroblast activity in presence of immune cells (NK-92in this case).^36^ We have shown with previous studies from our lab that fibroblast exposure to tumor cell-conditioned media induces stromal cell activation and secretion of a variety of cytokines.^26^ This activated phenotype is known to secrete IL-6, among other cytokines and chemokines. Hence, we also quantified fibroblast activated protein (FAP) by ELISA using media collected from CRC, CRC-HNLF, and HNLF-only TCs to evaluate presence of ‘activated’ fibroblasts.^26^ Next, we hypothesized that the inhibition of one of these cytokines would lead to increased NK cell homing to the TC. Here, we have achieved that via the IL-6 inhibitor Siltuximab. Following treatment with Siltuximab and with increasing dosage of Siltuximab, the combined action of NK cells and HNLF resulted in higher cytotoxicity in the tumor constructs compared to those samples without HNLF. This is shown starkly in the ATP assay (**Fig 5i and iii**). The accompanying LIVE/DEAD images also validated this result; most dead cells were visible in the tumor construct treated with the highest dose of Siltuximab in the samples containing both NK cells and fibroblasts compared to those which did not have fibroblasts (**Fig 5ii and iv**).

**Figure 5:**
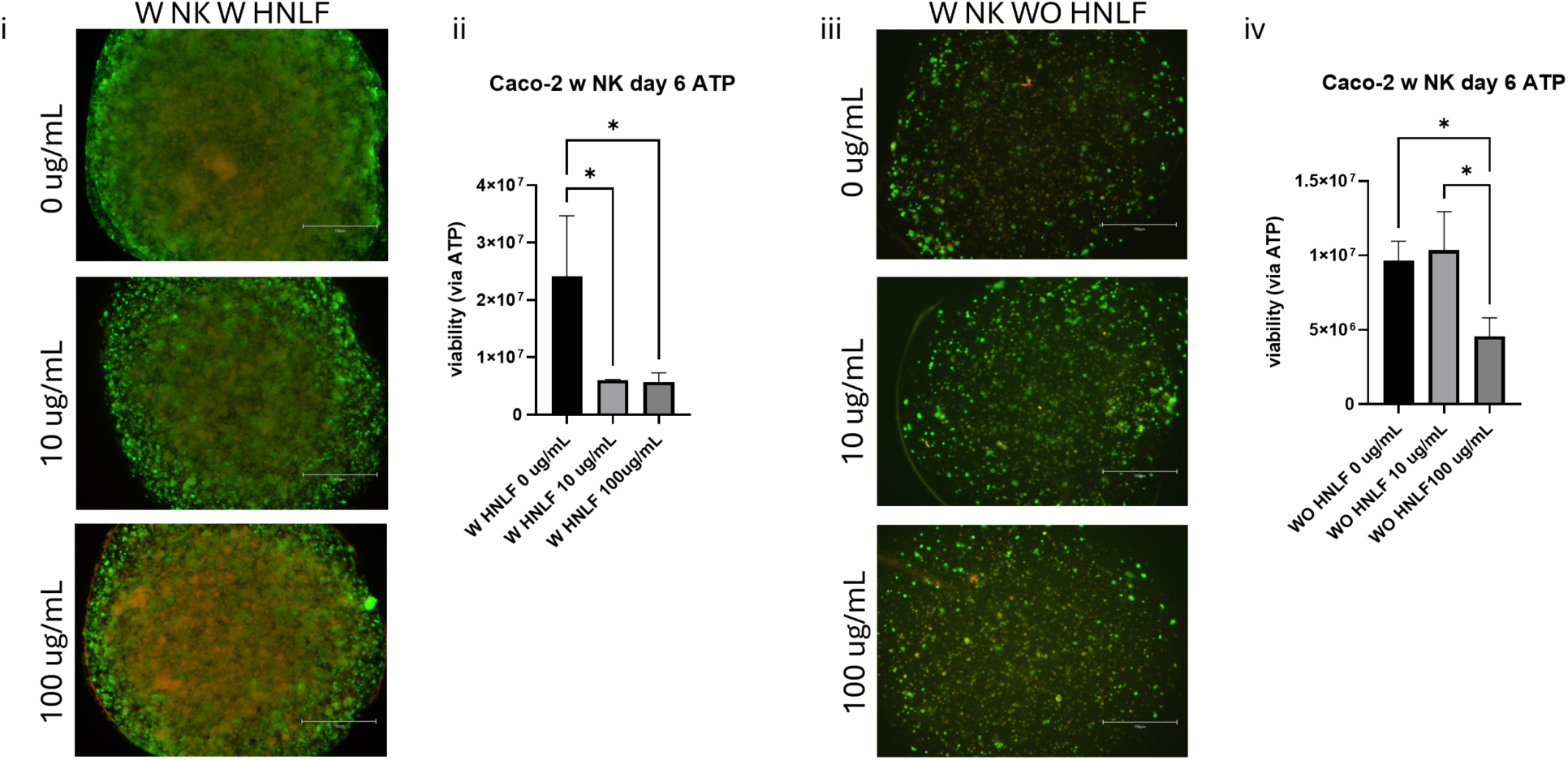
Viability studies with anti-IL-6 monoclonal antibody Siltuximab. (i) Representative images of LIVE/DEAD viability assay showed decreasing viability with increasing dose of Siltuximab in tumor constructs containing both NK cells and fibroblasts (ii) ATP assay confirmed this result as shown (iii) representative images of overall higher viability in samples with NK cells without fibroblast addition in tumor constructs with increasing dose of Siltuximab(iv) ATP assay confirmed this result as shown by the LIVE/DEAD images. Scale bar - 750 μm. Statistical significance: * p <0.05. Data are represented as mean + SD. Green - calcein AM-stained viable cells; Red - ethidium homodimer-1-stained dead cell nuclei.

### 2.5 Fibroblasts enable NK cells immune evasion in tumor cells via IL-6 in 3D microfluidic platform

Inspired by the results from the Transwell-based studies above, we wanted to test the influence of fibroblasts on NK cell functionality in a more advanced 3D bioengineered platform. Our lab has significant experience in designing and implementing microfluidic devices to model and study cancer – ranging from simple drug screening studies in tumor-on-a-chip systems to models of metastasis in our “metastasis-on-a-chip” platforms.^37–43^ This kind of a platform helps to ‘challenge’ NK cells to migrate towards and/or invade the tumor constructs in a more dynamic environment. Notably, microfluidic devices have been used extensively to study drug resistance, immune cell migration, and metastasis in wide variety of cancer-specific studies.^44–46^

Our team has an ongoing study in parallel to the study described herein, which tests the migratory capabilities of NK cells towards inhibitor drug-treated melanoma TCs and patient-derived organoids (PDOs) in a microfluidic device. Because we had this system up and running, we chose to deploy it here, but with and without fibroblasts in the TC component. While we recognize that this deviated from the CRC-focus thus far, our focus was more about the role of fibroblasts, regardless of the cancer type. Furthermore, understanding the role of fibroblasts in dictating the efficacy of NK therapy, and the potential importance of IL-6 – and other potential cytokines – in two cancers as opposed to one was an opportunity.

In this study, we fabricated tumor-on-a-chip (TOC) devices in which tumor constructs comprised of the same ECM-based hydrogels and tumor cells (A375 melanoma cells in this particular study) were located in chambers off to the side from a fluid channel that NK cells would be flown through. The logic for this design is that NK cells will need to sense chemokines or directly interact with the TC border, after which if the correct signals are present, they can bind and accumulate, followed by potential infiltration. (**Fig 6i-ii**) Red membrane dye DiI-labeled NK cells seemed to be migrating only to the drug-treated sample, particularly ABE, but also in the no drug control case (**Fig 6iii**). However, viability was higher in the control group as well as ABE compared to that seen for ALS (**Fig 6iv**). NK cell effector function was measured by the expression of Granzyme B and was significantly low in both ABE and the no drug-treated samples (**Fig 6v**). Also, cytokine levels for IL-6 and IL-8, which are often used to confirm fibroblast activity in presence of immune cells were higher in ALS-treated samples and these validated our hypothesis that fibroblasts were responsible in this case too, similar to the previously described studies detailing fibroblast involvement (**Fig 6iv-vii**).

**Figure 6:**
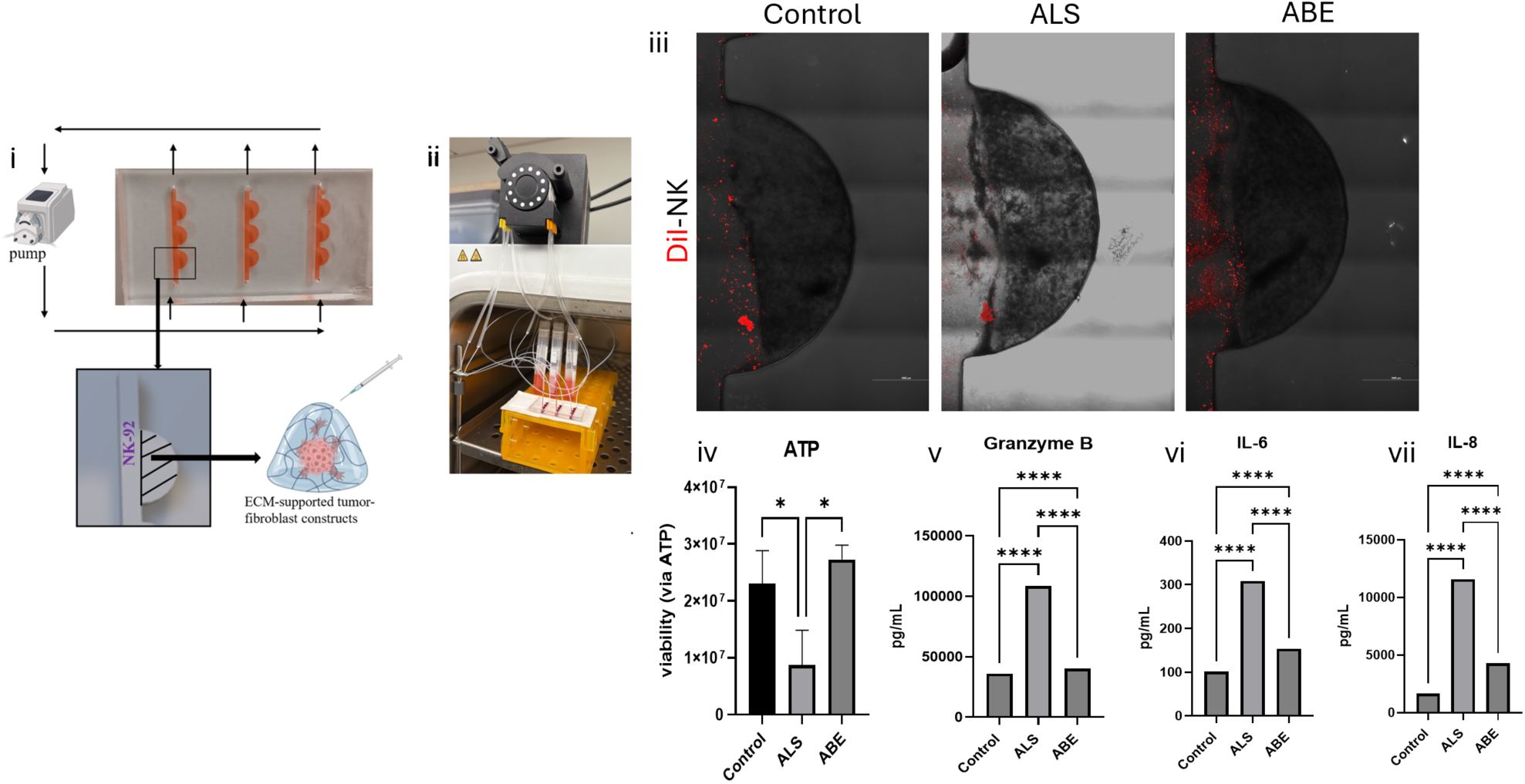
Schematic and general workflow for TOC device fabrication and NK cell migration, viability and chemokine profiles in TOC platform containing tumor cell-fibroblast constructs. (i) Schematic showing attachment of peristaltic pump to device, tumor cells and fibroblasts in chamber with NK cells in channel. (ii) TOC in operation: TCs are injected into finished device via insulin needle and media is flown via two-stop tubing connected to pump. (iii) Representative images of DiI-labeled NK cell migration showed more effective homing of NK cells to drug-treated particularly ABE-treated groups (iv) ATP assay however showed significantly higher viability in ABE-treated samples compared to ALS-treated samples. (v) NK cell effector function is validated via Granzyme B ELISA and showed significantly increased expression of this in ALS-treated samples (vi-vii) driver chemokine secretion profiles via IL-6 and IL-8 ELISAs showed significant expression of the same in the drug-treated samples, indicating fibroblast involvement and possible protective action of fibroblast towards tumor cells from the NK cells. Scale bars - 1000 μm. Statistical significance: * p < 0.05, ** p < 0.01, *** p < 0.001, **** p < 0.0001. Data are represented as mean +SD.

## 3. Materials and Methods

### 3.1 Cell passaging and culture

Highly aggressive HCT-116, mildly aggressive Caco-2, and epithelial SW480 cells were expanded in 2D on tissue culture plastic in 15 cm tissue-treated dishes with Dulbecco’s Modified Essential Medium (DMEM) until 80-90% confluence. Cells were then detached from the substrate with 0.25% Trypsin/EDTA and resuspended in growth media before use in further studies. Residual cells were cultured in DMEM, 5% FBS and 1% mixture of penicillin-streptomycin and L-glutamine until further use.

Human melanoma cells (A-375) were expanded in 2D on tissue culture plastic in 15 cm tissue-treated dishes with Dulbecco’s Modified Essential Medium (DMEM) until 80-90% confluence. Cells were then detached from the substrate with 0.25% Trypsin/EDTA and resuspended in growth media before use in further studies. Residual cells were cultured in DMEM, 5% FBS and 1% mixture of penicillin-streptomycin and L-glutamine until further use.

Human normal lung fibroblasts (HNLF) were expanded in 2D on tissue culture plastic in 15 cm tissue-treated dishes with fibroblast growth medium or FBM (FBM Basal Medium (CC-3131) and FGM-2 SingleQuots supplements (CC-4126), Lonza) until 80-90% confluence. Cells were then detached from the substrate with 0.25% Trypsin/EDTA and heat-inactivated fetal bovine serum. These were then resuspended in growth media before use in further studies. Residual cells were cultured in FBM complete media and/or cryopreserved until further use.

Human NK cells (NK-92) were expanded in untreated 2D tissue culture flasks (T25) with Alpha-MEM complete media. This media was aliquoted in 20 mL and modified with IL-2 added separately. This resulting media was replaced every 7 days to ensure fresh supply of IL-2 to the NK cells.

### 3.2 Tumor construct (TC) preparation

ECM-mimicking collagen-hyaluronic acid (HA) hydrogels were formed using methacrylated collagen (PhotoCol, Advanced Biomatrix, Carlsbad, CA) and thiolated HA (Glycosil, ESI-BIO, Alameda, CA). The HA was dissolved in water containing 0.1% w/v of the photoinitiator 4-(2-hydroxyethoxy) phenyl-(2-propyl) ketone to make 1% w/v solutions. Fifty-two μL of methacrylated collagen type I (Advanced BioMatrix, San Diego, CA) was added to 18.75 μL of the HA and 4.41 μL of neutralizing solution (for photo-collagen) in a small conical tube. Cells were added to this resulting solution and 5 μL TCs were constructed for each well of a 48-well plate previously coated with a thin layer of polydimethylsiloxane (PDMS), (**Fig 7**).

**Figure 7:**
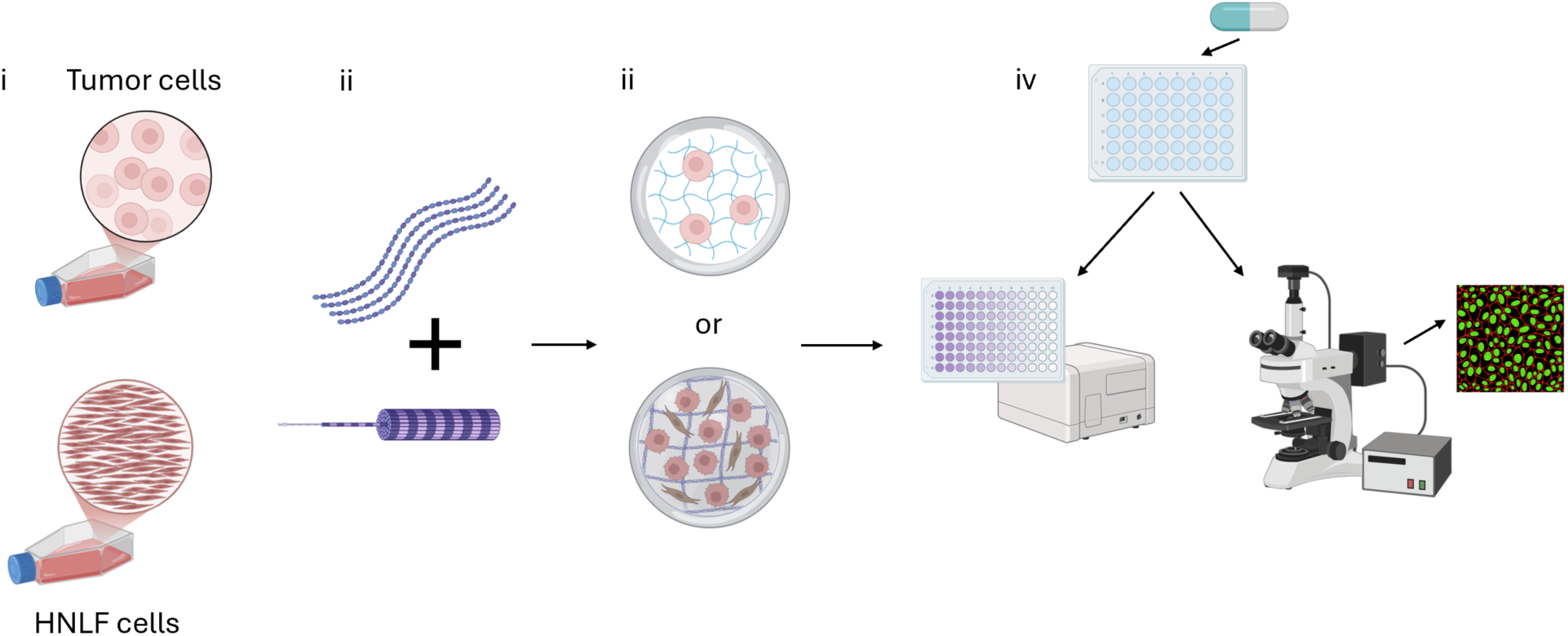
Schematic of experimental workflow. (i) Culturing of tumor cell lines and fibroblasts, (ii) Addition of hydrogel components like hyaluronic acid (top) and collagen (below), (iii) Formation of tumor cell constructs with and without fibroblasts, (iv) Viability assessment via luminescence-based ATP assay and LIVE/DEAD imaging after 24h chemotherapy.

### 3.3 Drug reconstitution and treatment

The colorectal cancer TCs were treated with several different drugs-Cisplatin (Sigma-Aldrich, St. Louis, MO), 5-Fluorouracil (Lifeline Cell Technology, Frederick, MD) and Regorafenib (Selleckchem/Selleck Chemicals LLC, Houston, TX)-for 24 hours. Concentrations of 10 μM and 100 μM were prepared from stock concentration of 10 mM for each drug. The drug-containing media was removed and replaced with fresh media after 24 hours. Viability assays were then done on days 1, 4 and 7 (d1, d4 and d7 respectively). Zero μM was chosen as the ‘no drug’ control for all conditions of all studies done here.

### 3.4 Phenotypic and functional assays

#### 3.4.1 Viability assays

LIVE/DEAD staining (LIVE/DEAD viability/cytotoxicity kit for mammalian cells; Thermo Fisher, Waltham, MA) is performed for every time point as mentioned above. Spent media was first aspirated from wells, after which 250 μL of mixture of PBS and DMEM (1:1) containing 0.5 μM calcein-AM and 2 μM ethidium homodimer-1 was introduced to each well. TCs were incubated for 30 minutes, after which fluorescent imaging was performed. Viability was also assessed via luminescence using Cell Titer-Glo 3D cell viability assay (Promega). Spent media was aspirated from wells containing the PTCs and 200 μL of 1:1 mixture of 3D ATP media and DMEM was added. The well-plate was gently shaken for 5 minutes and then incubated at room temperature for 25 minutes. These contents were then transferred to a Costar White Polystyrene 96 well Assay Plate (3912, Corning, NY) and luminescence was measured on a microplate reader (Varioskan LUX 3020-80022) using default settings for luminescence protocol.

#### 3.4.2 Fibroblast activation detection

Enzyme linked immunosorbent assays (ELISAs) were also performed to evaluate activated state of fibroblasts. Fibroblast activation protein (Abcam Human FAP ELISA kit-ab256404 kit) was used for this purpose. TCs were made using all three colorectal cancer cell lines, with and without fibroblasts. These were left untreated for 7 days after which media is saved for each condition. The standard protein concentrations were done in duplicates and samples in triplicates. Values were read using absorbance protocol (at 450 nm) on a microplate reader (Varioskan LUX 3020-80022).

#### 3.4.3 E-cadherin staining and imaging

TCs were made using Caco-2 cells, with and without the addition of fibroblasts. Growth media was added, and these were incubated for 7 days after which media was removed, and the TCs were fixed in 4% PFA overnight. After 24 hours, TCs were washed and stained for E-cadherin (CD324-E-Cadherin-Monoclonal Antibody-DECMA-1, Thermo Fisher, Waltham, MA) at the company recommended dilution.

#### 3.4.4 Imaging

LIVE/DEAD viability imaging was done on days 1, 4 and 7 using an EVOS M500 (Invitrogen) microscope. Appropriate filters were chosen for green and red fluorescence and imaging was done using 4x/10x objective. On the A1R confocal microscope (for E-cadherin staining), z-Stacks (200 μm) were obtained for each construct using filters appropriate for both red and green then superimposed.

#### 3.4.5 NK staining and flow

Fluorescently labeled NK-92 cells (Vybrant Multicolor Cell-Labeling Kit (DiO, DiI, DiD Solutions, 1 mL each), Invitrogen, Waltham, MA) were added via 0.4 µm pore sized cell culture inserts/transwells (Corning, NY) on day 6.

#### 3.4.6 Drug reconstitution and treatment

The TCs were treated with 1 μM Alisertib (ALS-Selleckchem 10 mM in 1 mL DMSO) and Abemaciclib (ABE-Selleckchem 5 mg) respectively, for 5 days. Both the drugs were reconstituted from their stock concentration of 10 mM to 1 μM. Zero μM was used as no drug control/vehicle for all conditions. Output metrices were performed on day 6.

#### 3.4.7 Microfluidic device fabrication

Microfluidic device molds were designed using AutoCAD. Molds were printed using the H-Series 3D printer (CADworks 3D, Concord, ON, CA) with CADworks 3D Mastermold Resin and were used to produce devices out of polydimethylsiloxane (PDMS, Sylgard 184, DOW Chemical). The PDMS devices were left in the oven overnight to solidify and then bonded to glass slides via plasma treatment. There were inlet and outlet points in every channel for media addition, punched using 5 mm biopsy punches. Orange-yellow two-stop tubing (0.51 mm) was attached to these channels and then to a six-channel pump (Elemental Scientific MP^2^ Stand Alone Precision Micro Peristaltic Pump) to ensure flow of media. The velocity of flow was set at 0.8 μL/sec and 5 rpm. Hydrogel-encapsulated TCs were injected into the device chambers using a small (0.31×8mm) insulin gauge syringe and then photopolymerized in place using a brief UV light pulse (365 nm, 18 W cm−2 for 2s to initiate crosslinking of the hydrogel.

#### 3.4.8 Inhibition study

The IL-6 inhibitor Siltuximab (Selleckchem-anti IL-6 CNTO 328) was used. Ten μg/mL and 100 μg/mL concentrations were prepared from 1 mg stock concentration while 0 μg/mL was used as no drug control/vehicle.

#### 3.4.9 Protein detection assays and cytokine profile identification

Spent media from TCs not treated with any drugs and without any NK cell addition was collected and used to detect presence of chemokines specifically CCL5 (Human RANTES ELISA Kit-ab174446), CXCL10 (Human IP-10 ELISA kit-ab83700), IL-8 (Human IL-8 ELISA kit-ab214030) and IL-6 (Human IL-6 ELISA kit-ab178013) via enzyme-linked immunosorbent assays (ELISA).

#### 3.4.10 NK cell cytotoxicity validation

Spent media was collected from TC samples on day 6 and Granzyme B (Human Granzyme B ELISA kit ab235635), TNF-α (Human TNF alpha ELISA kit ab181421) and IFN-γ (Human Interferon gamma ELISA kit ab300323) ELISAs were performed to evaluate NK cell effector function.

#### 3.4.11 Statistical analysis

All samples/groups were maintained in triplicates. These triplicate values for each condition were averaged and analyzed via using Graph Pad Prism (GraphPad, La Jolla, CA) software. ANOVA followed by Tukey’s was used to determine statistical significance. P-value less than 0.05 was considered significant.

## 4. Discussion & Conclusion

The communication between cells in the TME is crucial to understand how to design effective therapeutic options. Here, we studied how fibroblasts influence therapies in ECM hydrogel-supported 3D tumor construct models, with our work focusing in on the NK cell-based immunotherapy. We observed that the fibroblasts appeared to shield the CRC tumor cells from chemotherapy and also reduced effective NK cell effector function following kinase inhibitor intervention meant to aid in driving NK cell effector function. We found that fibroblasts play a tumor-aiding role by secreting different chemokines, most notably IL-6. We also noted similar patterns of fibroblast activity in a dynamic microfluidic device containing 3D melanoma constructs that were treated with NK cells.

Our initial focus was on how fibroblasts impact chemotherapy in CRC. As we show in **Fig 1** using Caco2 TCs, there are some trends that arise, suggesting that chemotherapies might be less effective in the presence of fibroblasts. This is not a new finding, but using our bioengineered in vitro platform, we had hoped that we could observe a clear impact of fibroblasts one way or another.

The inconsistency shown in **Fig 1** was even more pronounced in the ATP assay for the Caco2 studies, and both LIVE/DEAD and ATP assays for SW480 and HCT-116 cells (**Fig S1, S2, S3, S4, and S5**). As a result, while the differences due to fibroblasts were inconsistent, they provided motivation for continued study, but in a different therapeutic context. We turned our attention towards immunotherapy; specifically, NK cell-based immunotherapy.

Studies have shown how secretion of cytokines such as IL-6 help CAFs to modulate immune cells to generate an immunosuppressive TME through different pathways including reducing thus effector function of NK cells.^47^ It is well established that there is ample dialogue between CAFs and NK cells in the TME in various malignancies. Inflammatory CAFs can produce IL-6 which impedes NK cell function suppression by limiting IFN-γ secretion.^48^ Interestingly, in gastric tumors, CAFs can promote iron overload and iron-dependent lipid peroxides which result in impairment of NK cell effector functions via ferroptosis. Hence, in gastric cancer, efficient NK cell activity is inversely related to the presence of CAFs, and this could be the case in other cancers as well.^49^

A signaling pathway that we identified as important was IL-6 signaling. The IL-6/STAT3 signaling pathway is crucial in the development of colorectal cancer. IL-6 levels are usually upregulated in CRC patients, and this is associated with patient prognosis-IL-6 expression is higher in CRC tissue samples than in normal tissue samples. IL-6 activates JAK/STAT3 which in turn drives malignant pathways in tumor cells which influence tumor cell invasion, metastasis, and angiogenesis. IL-6 can further function as a paracrine cytokine to also facilitate CRC cell proliferation as well as oppose tumor cell apoptosis. IL-6 is an important tumor promoting factor in various other cancers including melanoma, glioma, lymphoma, and others. IL-6 overexpression is also linked to chronic inflammation leading to inflammatory bowel disease and eventually tumor formations.^50^ CAFs via IL-6 can induce chemoresistance of gastric cancer cells. This is achieved via STAT3-pathway which lowers apoptosis of tumor cells resulting from chemotherapy. Thus, a poor response to chemotherapy is usually associated with IL-6 expression.^51^ While somewhat inconsistent, in some CRC cell lines with some chemotherapy agents, we observed decreased tumor cell killing. Fibroblast-based or -induced IL-6 production could potentially be involved in these instances. We did not inhibit IL-6 in our chemotherapy studies given the inconsistent responses we observed with and without fibroblasts present. However, in future studies we are interested in down-selecting to the chemotherapeutic agents that were impacted by the presence of fibroblasts and performing IL-6 inhibition studies like those we performed in our NK cell-based studies.

With respect to NK cells, IL-6 has been documented to help mobilize effector NK cells towards some tumor cells, but it has also been shown to help other tumor cells escape immune surveillance and clearance. This IL-6 enabled hindrance of normal NK cell cytotoxic functions can be achieved in various ways.^52^ IL-6 is associated with increased PD-L1 expression on tumor cells, which can lead to NK cell exhaustion through the immune checkpoint PD-1/PD-L1 interaction. Elevated IL-6 in the TME can further promote the differentiation of helper T cells into Th17 and regulatory T cells which can indirectly suppress NK cell activity and favor tumor growth. Also, IL-6 can enhance cancer stem cell-like properties in CRC cells, helping them resist NK cells.^53^ In our systems, we of course did not have T cells present, but even cancer stem cells can have subpopulations that could be or could be induced to be more akin to cancer stem cells. Future studies that that evaluate the impact of IL-6 on cancer stem cell gene and protein expression profiles will some next steps that we will explore.

Finally, we report a previously unreported mechanism of fibroblast impairment of NK cell function via IL-6. However, further studies are needed to confirm NK cell effector function as well as identify activated fibroblasts state to better describe this crosstalk between immune and stromal components. There is also a scope to look at which other pathways such as IL-8 which is documented to be often used by fibroblasts to aid tumor progression^54–58^ and interfere with the normal NK cell cytotoxic functions in different malignancies.^59–61^

In terms of therapeutic intervention, we utilized the anti-IL-6 monoclonal antibody Siltuximab. This drug was approved for cases of human immunodeficiency virus negative and human herpesvirus-8 negative multicentric Castleman’s disease by the US FDA in 2014 with a dose of 11 mg/kg over 1-hour intravenous infusion every 21 days. Interestingly, an advanced phase clinical study aimed at highlighting the pharmacokinetics of the same drug as monotherapy in CRC did not yield significant results even though no side-effects were documented.^62^ Despite the lack of significant anti-cancer results, this result paired with the results described here might suggest that Siltuximab on its own is clearly not an effective anti-cancer treatment. But could it prime the TME to make that TME more susceptible to an immunotherapy, such as NK cellular therapy? Our goal is to further explore this NK cell - IL-6 – fibroblast connection and Siltuximab inhibition in more advanced human cell-based *in vitro* tumor- and organ-on-a-chip models, including using patient-derived tumor construct models.

## Acknowledgements

We acknowledge resources from the Campus Microscopy and Imaging Facility (CMIF) and the OSU Comprehensive Cancer Center (OSUCCC), The Ohio State University. This facility is supported in part by grant P30 CA016058, National Cancer Institute, Bethesda, MD. A Skardal acknowledges support from the OSUCCC and the OSUCCC Translational Therapeutics Pilot Award Program.

## Supplemental materials

### Supplementary information

Higher viability and denser growth were seen in the 3D HCT-HNLF TCs compared to HCT only TCs (**Fig S1iii**) treated with Cisplatin. Cisplatin inhibits DNA replication and transcription via formation of cross-links, leading to cancer cell death. This resistance is brought about by alterations in DNA damage signaling as well as defects in the cellular mismatch repair pathway.^64^ However, colorectal cancer cells can form resistance to cisplatin, making it a less effective therapy in CRC. In our models as well, overall viability was greater in HCT-HCT-HNLF, and SW480-SW480-HNLF (**Fig S1ii**) samples compared to all Caco-2 TCs (**Fig S1i**). Additionally, Cisplatin when used with other drugs such as 5-FU holds the potential to be more effective.

**Figure S1:**
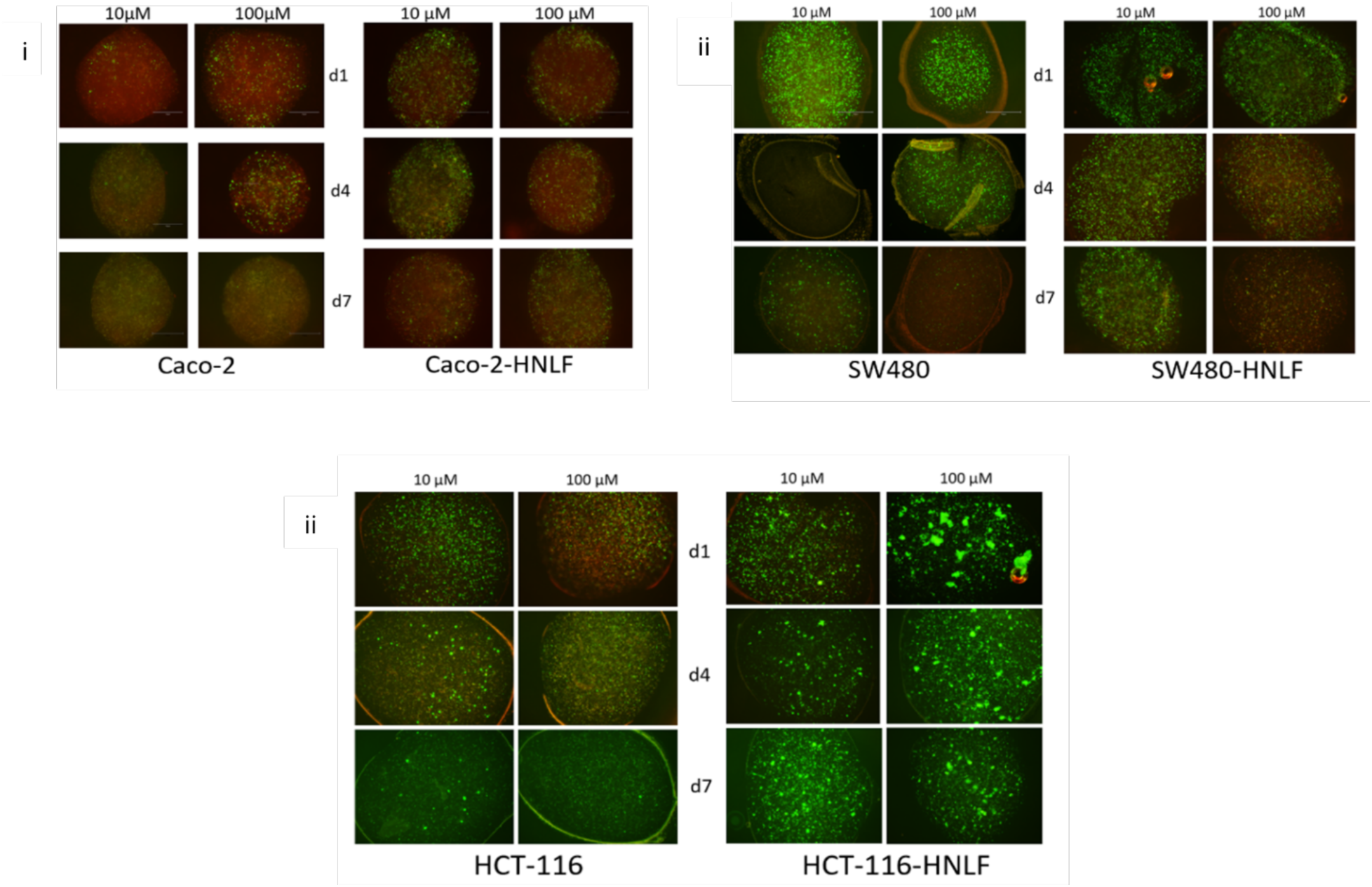
Post Cisplatin treatment-viability in 3D hydrogel-supported colorectal cancer tumor constructs made with and without fibroblasts via LIVE/DEAD assay on days 1,4, and 7. (i) Representative imaging of LIVE/DEAD viability assay in Cisplatin-treated Caco-2 cells with and without fibroblasts in 3D tumor constructs showed lowered viability in both 10 and 100 μM treated samples (ii) There was not as much difference in viability between tumor cell only and tumor cell-fibroblast samples for SW480 cells, although the cellular growth is visibly higher than Caco-2 samples. (iii) Viability is comparatively higher in HCT-116-HNLF tumor constructs compared to HCT-116 only samples, and overall higher among the models of the three different cell types. Scale bar - 300 μm. Green—calcein AM-stained viable cells; Red—ethidium homodimer-1-stained dead cell nuclei.

Also, upon Regorafenib treatment, the viability was higher in SW480-HNLF compared to SW480 TCs only (**Fig S2ii**). Regorafenib is a multi-kinase (receptor tyrosine kinases such as KIT and RET) inhibitor used in metastatic CRC where it works on proteins involved in cancer growth, angiogenesis and the tumor microenvironment in general. With respect to the HCT samples, there were also a smaller number of dead cells overall and more confluent live cells (**Fig S2iii**). The overall viability in the Caco-2 and Caco-2-HNLF samples appeared to be the lowest with more dead cells visible at the center of the constructs (**Fig S2i**). Now, Regorafenib is a small molecule multi-kinase inhibitor. It has been used in multiple cases as treatment for CRC but in cases where there has been some sort of previous therapeutic intervention. That could be a reason why the responses were not uniformly effective in our models across the different conditions.^65,66^

**Figure S2:**
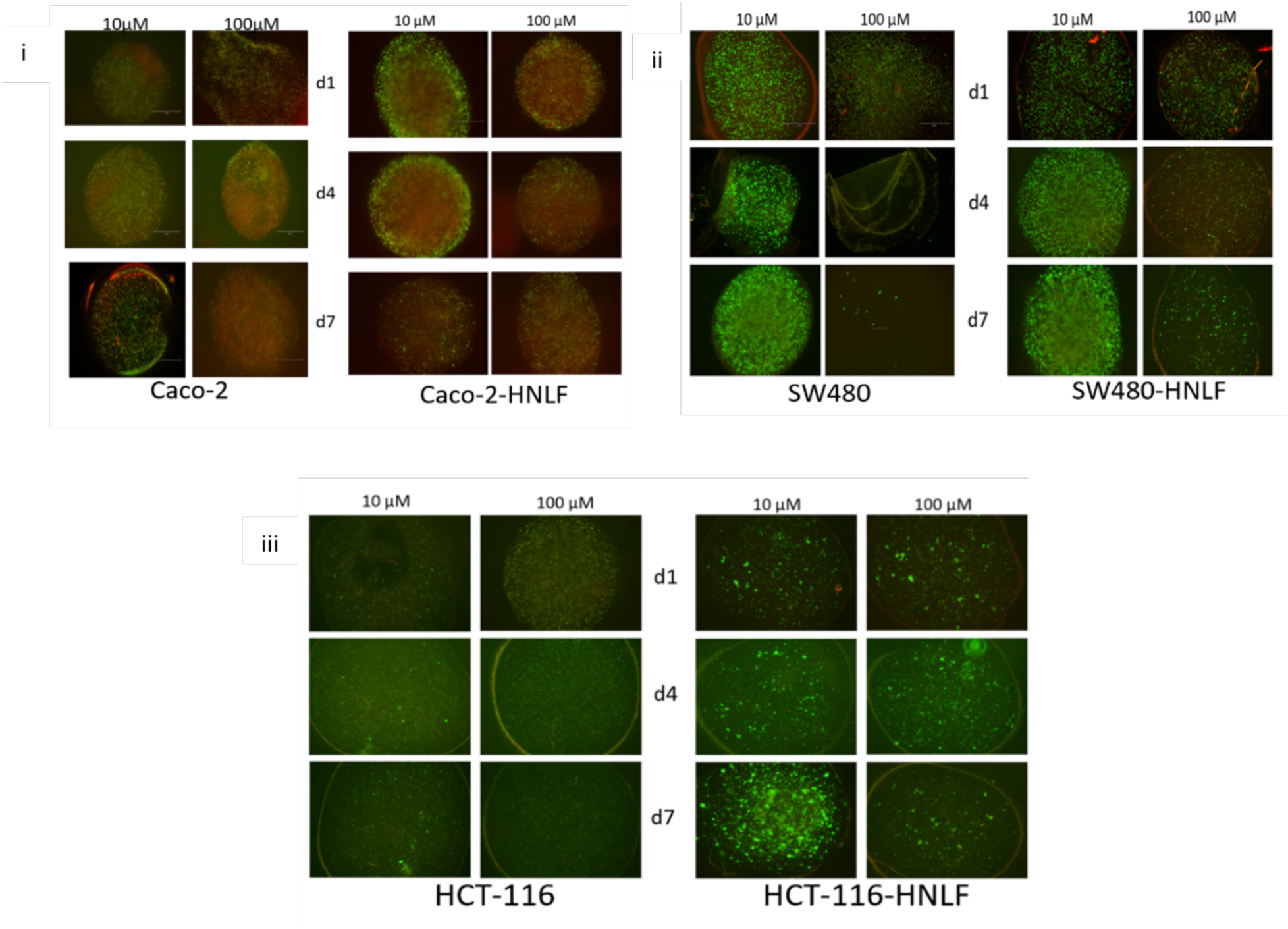
Post Regorafenib treatment-viability in 3D hydrogel-supported colorectal cancer tumor constructs made with and without fibroblasts via LIVE/DEAD assay on days 1,4, and 7. (iii) After Regorafenib treatment, all Caco-2-containing samples show more dead cells particularly at the center of the constructs. (ii) The viability is higher in SW480-HNLF constructs than in SW480 only samples (iii) The same is true for the HCT samples, but the constructs are not as dense and finally (iv) Scale bar - 300 μm. Green—calcein AM-stained viable cells; Red— ethidium homodimer-1-stained dead cell nuclei.

**Figure S3:**
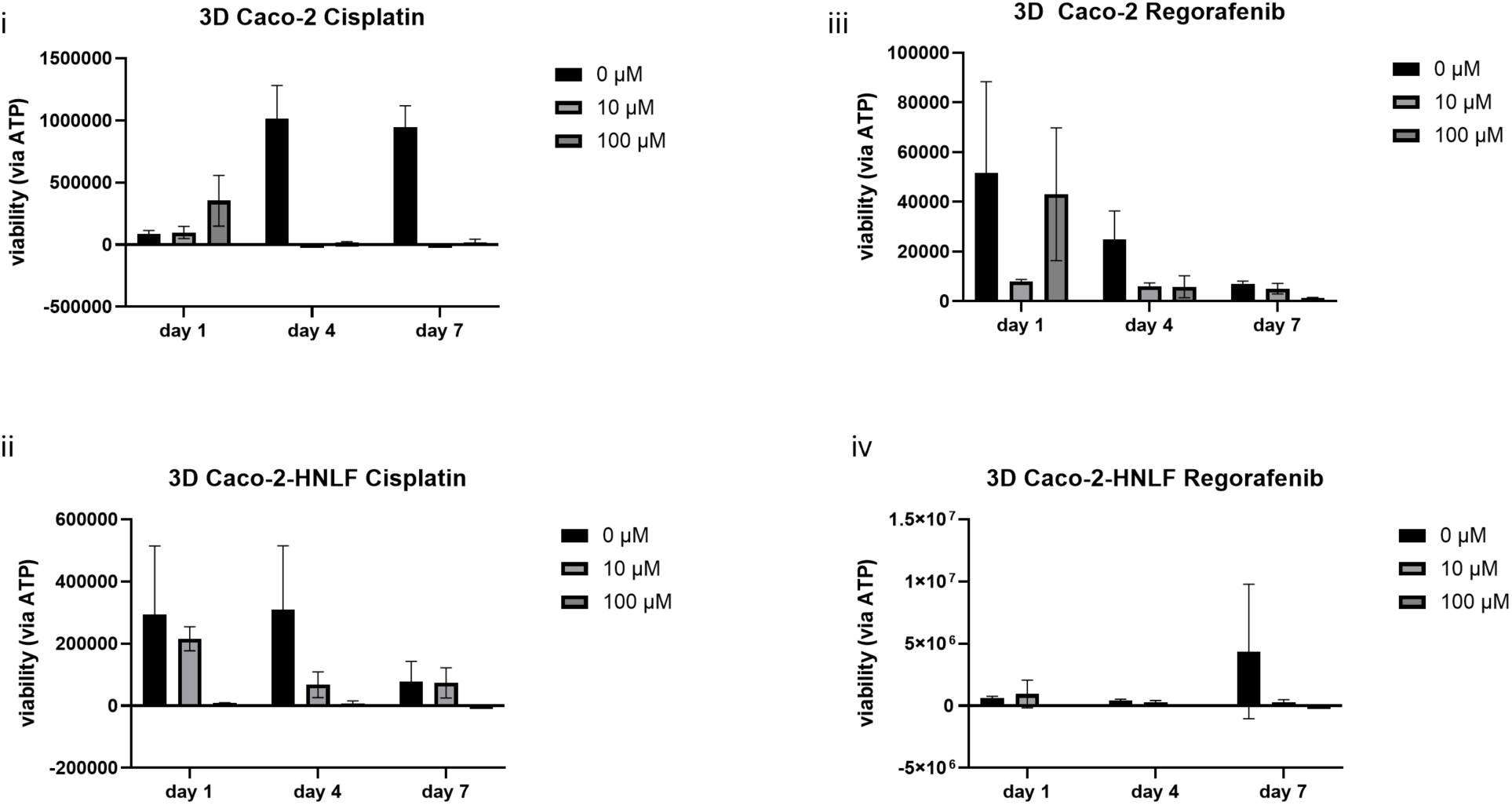
Viability measurements of Caco-2 and Caco-2-HNLF tumor constructs via ATP assay after treatment with Cisplatin and Regorafenib. (i-ii) Luminescence-based ATP assay showed reduced viability in Caco-2-only constructs when compared with Caco-2-HNLF constructs specifically with respect to 10 μM Cisplatin treatment. (iii-iv) This was not visible for the TCs treated with Regorafenib for any specific concentration of drug.

**Figure S4:**
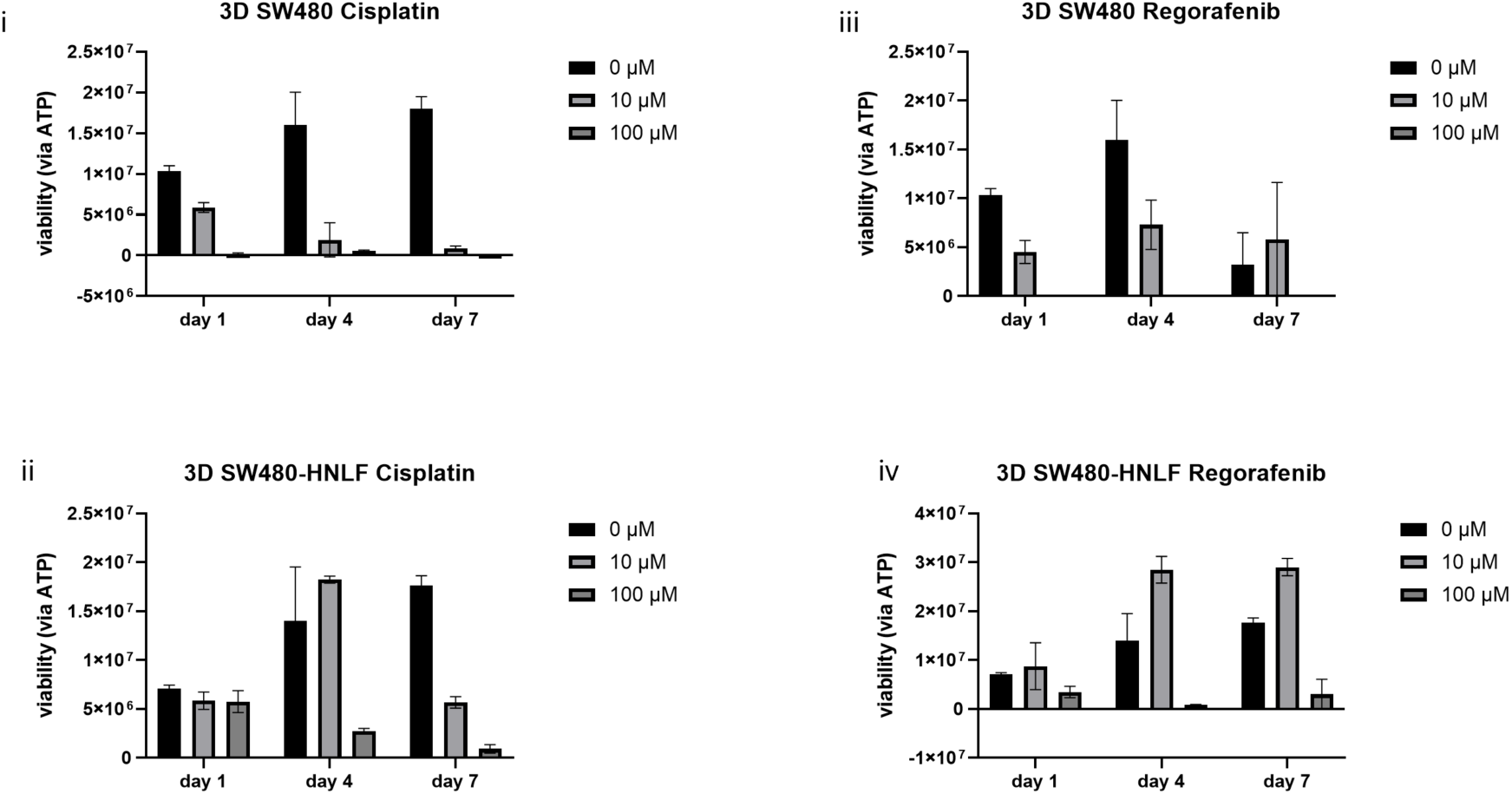
Viability measurements of SW480 and SW480-HNLF tumor constructs via ATP assay after treatment with Cisplatin and Regorafenib. (i-ii) Luminescence-based ATP assay showed reduced viability in SW480-only constructs when compared with SW480-HNLF constructs with respect to 10 μM as well as 100 μM Cisplatin treatment. (iii-iv) This was also visible for the TCs treated with Regorafenib for the same concentrations.

**Figure S5:**
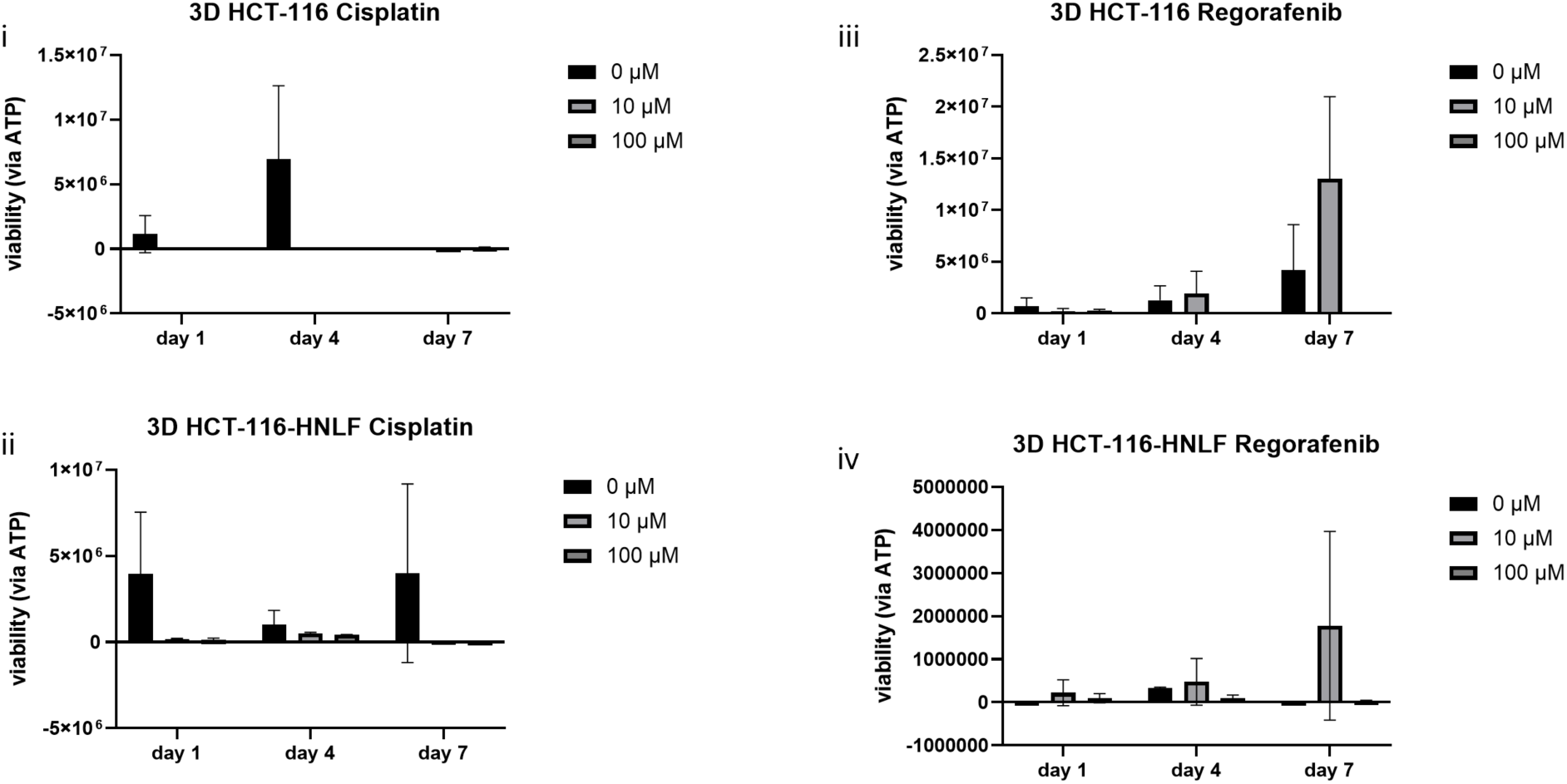
Viability measurements of HCT-116 and HCT-116-HNLF tumor constructs via ATP assay after treatment with Cisplatin and Regorafenib. (i-ii) Luminescence-based ATP assay showed reduced viability in HCT-116-only constructs when compared with HCT-116-HNLF constructs with respect to the day 4 10 μM as well as 100 μM Cisplatin treatment. (iii-iv) This was not visible for the TCs treated with Regorafenib for any of the drug concentrations used.

